# Cell volume distributions in exponentially growing populations

**DOI:** 10.1101/673442

**Authors:** Pavol Bokes, Abhyudai Singh

## Abstract

Stochastic effects in cell growth and division drive variability in cellular volumes both at the single-cell level and at the level of growing cell populations. Here we consider a simple and tractable model in which cell volumes grow exponentially, cell division is symmetric, and its rate is volume-dependent. Consistently with previous observations, the model is shown to sustain oscillatory behaviour with alternating phases of slow and fast growth. Exact simulation algorithms and large-time asymptotics are developed and cross-validated for the single-cell and whole-population formulations of the model. The two formulations are shown to provide similar results during the phases of slow growth, but differ during the fast-growth phases. Specifically, the single-cell formulation systematically underestimates the proportion of small cells. More generally, our results suggest that measurable characteristics of cells may follow different distributions depending on whether a single-cell lineage or an entire population is considered.

## 1 Introduction

Each living cell is an individual entity occupying a given volume enclosed by the cell membrane [1]. Homeostatis of cell volume is due to balance between cell growth and division. Growth in cell volume is understood to occur continuously in time and is often assumed to be exponential. Cell division is typically represented as a discrete event at which the volume of a mother cell abruptly changes into the volume of either daughter cell [19]. Specifically, symmetric division means that each daughter obtains exactly one half of their mother’s volume. The contents of a mother cell, including its transcriptome and proteome, are also divided between its daughters. Fluctuations in cell volume due to cell growth and division can therefore correlate with gene-expression noise [3].

There are (at least) two alternative approaches to the modelling of cell-growth dynamics. In the first approach, one follows a single cell line, discarding the other daughter cell at each division. In the single-cell approach, the time-dependent cell volume can be represented by a piecewise deterministic [6] or drift-jump Markovian process [15]. One is interested in the probability distribution of the random process, in particular at steady state. In the second approach, one follows both daughter cells, and is interested in the dynamics of the population size as well as the distribution of cell volumes among the population, in particular in the large-time limit. A question of interest is whether the probability distribution obtained from the single-cell approach and the population distribution obtained from the population approach are the same or different. The difference between single-cell and population approaches can be relevant in a number of applications, e.g. in cancer biology, which allow for experimental setups in which the reproductive history of a cell can be traced [10]. We will examine this problem for a particular type of volume-growth model.

The maintenance of homeostasis requires that cells actively control their proliferation [14, 16, 18]. The necessary feedback can be exerted through e.g. the cell’s age [9], its current size [13], the size at its inception [2], or by a combination of these mechanisms [11, 17]. In this manuscript we specifically focus on size-based regulation. Within the framework of a relatively simple model, we will proceed towards the following goals: (i) develop exact and efficient stochastic algorithms to simulate the volume growth process; (ii) characterise the large-time asymptotic behaviour of the process by formulating and solving a master equation; and (iii) draw conclusions about the similarities and differences between the single-cell and the whole-population approach to modelling cell growth.

The outline of the paper is as follows: in Section 2 we introduce and logarithmically transform the model. In Section 3 we present an iterative algorithm for simulating the single-cell version of the model and a recursive algorithm for the simulation of the whole-population version. In Section 4 we introduce the concept of periodicity in the context of the current model. In Sections 5 and 6 we formulate the master equation and provide tractable closed-form formulae for large-time solutions in the single-cell and whole-population cases. In Section 7 we present the main implications of the current work on the dynamics of cell growth and cell volume distributions. In Section 8 we extract conclusions from the presented analysis.

## 2 Model fundamentals

Our model for cell-volume growth is based on the following fundamentals:

1. Cell volume grows exponentially in time. Specifically,

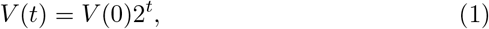

where *t* is the time since birth. Time is measured in units of the volume doubling time.
2. A mother cell can divide into two daughter cells. Either daughter cells obtains one half of the mother’s volume. From a single-cell viewpoint, at the time of division the volume changes abruptly according to the mapping

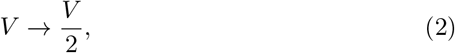

where the volume on the left-hand side represents the volume of the mother right before the division and a daughter’s volume is on the right-hand side.
3. Cells divide with a volume dependent stochastic rate γ(*V*). We specifically focus on the case of

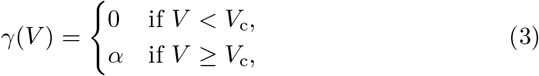

where *α* is a constant rate and *V*_c_ is a critical volume threshold.

It turns out that it is much more convenient to use a logarithmic transformation of cell volume defined by

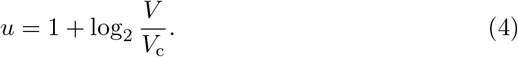

By (3), cells cannot divide before they reach the critical volume *V*_c_. Hence the cell volume is always greater than 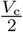, and the log-volume, as defined by (4), is always positive.

The model fundamentals (1)-(3), when expressed in the language of log-volume, read as follows:

1. Dividing (1) by the critical volume *V*_c_ and taking the binary logarithm gives

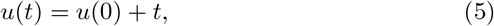

i.e between divisions a cell’s log-volume grows linearly with unit rate.
2. Taking the binary logarithm of (2) divided by *V*_c_ gives

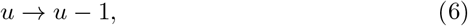

meaning that, upon division, a cell’s log-volume decreases by one.
3. Since requiring *V* ≥ *V*_c_ is equivalent to *u* ≥ 1, the dependence (3) of the stochastic rate on volume translates into

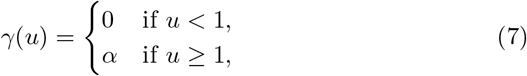

in terms of the log-volume.

Table 1 summarises the model fundamentals in the linear and logarithmic volume scales.

**Table 1.** Model formulation in terms of Volume *V* (left column) and Log-Volume 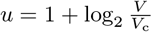 (right column).

| | Volume $V$ | Log-Volume<br>($U = 1 + \log_2 \frac{V}{V_c}$) |
| --- | --- | --- |
| Growth | $V(t) = V(0)2^t$ | $u(t) = u(0) + t$ |
| Division Map | $V \rightarrow \frac{V}{2}$ | $u \rightarrow u - 1$ |
| Division Rate | $\gamma(V) = \begin{cases} 0 & \text{if } V < V_c, \\ \alpha & \text{if } V \geq V_c, \end{cases}$ | $\gamma(u) = \begin{cases} 0 & \text{if } u < 1, \\ \alpha & \text{if } u \geq 1, \end{cases}$ |

## 3 Stochastic simulation

In this section we present an algorithmic approach the model of cell-volume growth based on the fundamentals presented in Section 2. Hereby we distinguish two versions of the model.

**Single-cell version** Upon cell division, one of the daughter cells is followed, the other discarded. We are interested in the probabilistic description of the cell volume along an arbitrarily chosen lineage.
**Population version** Both daughter cells are followed upon cell division. We are interested in the growth of the number of offspring and in the distribution of volume across the population.

Throughout this section we will operate with log-volumes of cells rather than their volumes (see Section 2 for explanation).

A common building block in both versions of the algorithm is to sample the waiting time *τ* for division of a cell which currently has log-volume *u*. The waiting time consists of two parts: the deterministic time required to reach the critical log-volume of one; the stochastic time that it takes to divide once the critical threshold has been passed. Since log-volume grows with unit rate, the deterministic waiting time is 1 – *u* if u < 1 and zero if it is greater than one. After crossing the threshold division has a constant propensity *α* to occur; it follows that the stochastic waiting time will be exponentially distributed with mean 1/α. Putting the deterministic and stochastic parts of the waiting time *τ* together, we find that it can be sampled as

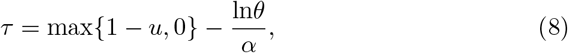

where *θ* is drawn from the uniform distribution on the unit interval. We have thereby used the well known fact that −ln*θ* is exponentially distributed with unit mean.

We are now ready to go through the individual steps of the single-cell version of the simulation algorithm (Algorithm 1). The algorithm requires the following inputs: the model parameter *α* which gives the (post-threshold) division rate; the initial log-volume *u*_0_; the time points *t*_0_, *t*_1_, …, *t_m_* at which we wish the log-volume to be recorded. We assume that these time points are ordered from the lowest to the largest. Time *t*_0_ is understood to be the initial time at which the log-volume is given by the intial value *u*_0_. The algorithm returns the sampled value *u*_1_, …, *u_m_* of log-volumes at the given time points.

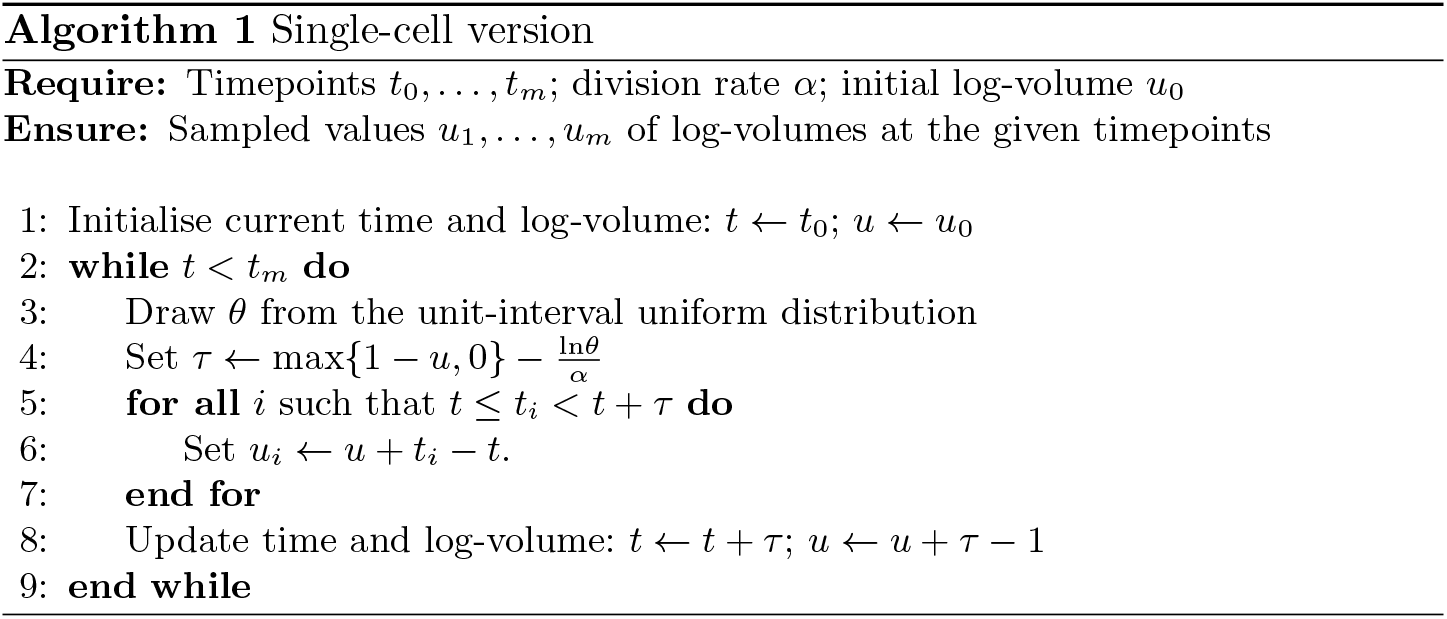

The algorithm starts by initialising the current time *t* and log-volume *u* with the initial values *t*_0_ and *u*_0_ (Line 1 in Algorithm 1). The next steps are repeated while the current time *t* is less that the largest time point *t_m_* at which a recording of the log-volume is sought (Lines 2-9): first, the waiting time until the next division is sampled in Lines 3-4 using the formula (8); second, log-volumes are recorded at all recording times *t_i_* which fall between the current time *t* and the time *t* + *τ* of next division (Lines 5-7); third, the current time and log-volume are updated to the post division values (Line 8).

We are now well positioned to proceed to the population version of the simulation algorithm (Algorithm 2). The population version is only marginally more elaborate than the single-cell version thanks to the use of recursion. Algorithm 2 requires the same input as Algorithm 1, but provides a different output, returning for each given time point a list of log-volumes across the whole population. By a list we understand a collection of elements (here log-volumes), some of which may be present in the list multiple times. We may append a number to a list; we may query how many times a given element is present in the list; we may query the total number of elements in the list — here the population size.

The algorithm proceeds as follows. First, we make sure that the lists are empty initially (Lines 1-3). Then we make a call to the procedure CELL (Line 15). The CELL procedure is defined recursively in Lines 4-14. The procedure calculates the contribution made by a cell that is introduced into the population at time *t*_0_ with log-volume *u*_0_, and by the entire offspring of that cell, to the lists of log-volume recordings. The cell’s individual contribution is calculated in Lines 5-9. Comparing Lines 5-9 in Algorithm 2 to the corresponding passage in Algorithm 1 (Lines 3-7), we note that the two sections of code differ only in that the cell’s log-volume is either stored as a single value (Algorithm 1, Line 6) or added to a list potentially containing multiple values (Algorithm 2, Line 8). The contribution of the cell’s offspring to the log-volume recordings is calculated in Lines 11-12 by making a recursive call to the CELL procedure for either of its daughter cells. Cells that are born after the last recording time point *t_m_* cannot make contribution to the log-volumes recordings. For this reason, the recursive calls are made only if the mother cell divides before the time *t_m_* of last recording (Line 10). In this manner, we make sure that the recursion does not continue *ad infinitum*.

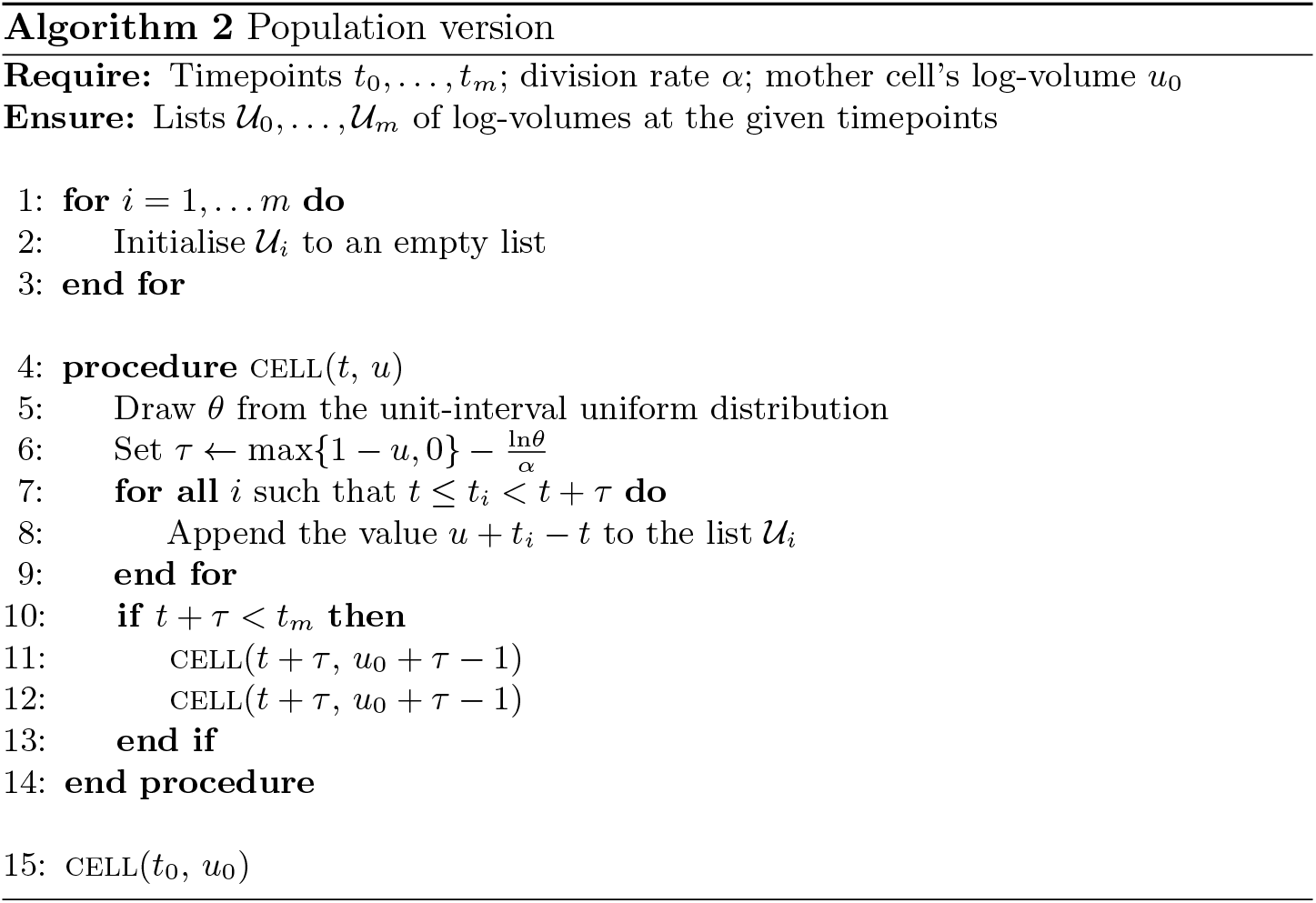

## 4 Periodicity

Regardless of the particulars of the growth-control mechanism, a minimalistic model based on exponential growth and symmetric division, which we shall consider here, exhibits a type of periodic behaviour [4, 5, 7]. Specifically, the volume, measured in units of its doubling time, of a daughter cell at time *t* > 0 is equal to the volume of the mother cell at time *t* = 0 multiplied by 2^*t*–*n*^, where *n* is the daughter cell’s generation. A couple of important observation follow immediately from this. First, possible cell volumes are restricted to a discrete set of values at any given time. Second, cell-volume measurements taken at different times cannot be equal unless the times of measurement differ by an integer multiple of the volume doubling time. We will see in what follows that this periodicity with respect to the volume doubling time has important consequences for the cell growth process that persist even in the asymptotic limit of large times.

Let *t*_0_ be the initial time and *u*_0_ be the log-volume of the mother cell at the initial time. These two are input values in stochastic simulation. At time *t* > *t*_0_, the log-volume *u*(*t*) of a daughter cell is constrained to the discrete set

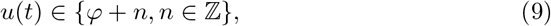

where 0 ≤ *φ* < 1 is a phase defined by

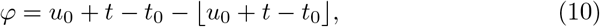

where ⌊*a*⌋ denotes the floor of *a* (the nearest integer lower than the real number *a*). The choice of the value of *n* within the discrete set (9) depends inversely on the number of divisions of the first mother cell up to the daughter.

The constraint (9) holds regardless whether the single-cell or the population approach to modelling cell-volume dynamics is taken. Additionally, it holds regardless of the choice of the log-volume-dependent division rate *γ*(*u*). Specifically, for the threshold-like dependence (7) we know that the log-volume has to be positive, implying that the *n* in the constraint (9) has to be non-negative.

The presence of the constraint (9) is best explained graphically (Figure 1, top panel). While the cell does not divide, its log-volume increases along a straight line with unit slope. When it divides, the log-volume transfers to a parallel line with intercept one unit lower. Regardless of the timing of cell divisions, the log-volume trajectories are constrained to a discrete union of parallel lines whose intercepts differ by an integer. The constraint (9) is obtained by taking a cross-section at time *t* of these parallel lines.

**Fig. 1.**
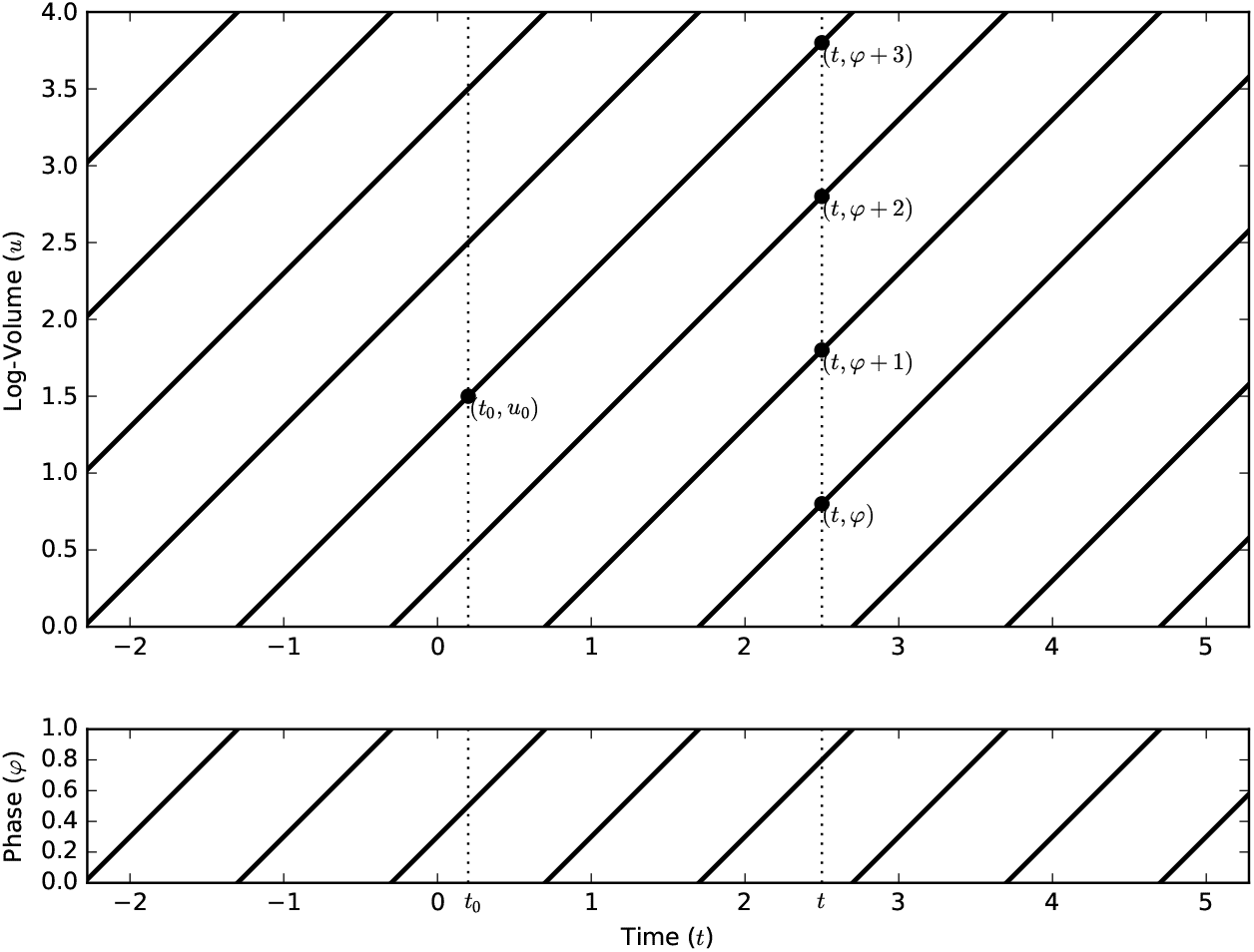
Periodicity of the cell-volume process. At any given time, the log-volume belongs to a discrete constraint set (top panel). Which of the constraint sets applies is determined by the phase *φ*, which is a one-periodic function of time *t* (bottom panel).

The constraint sets (9) are parametrised by the phase *φ*, which is a one-periodic function of time *t* (Figure 1, bottom panel). The log-volume *u*(*t*) visits each constraint set periodically with unit period. Different phases give disjoint constraint sets. The union of all constraint sets over phases 0 ≤ *φ* < 1 gives the entire state space of real log-volumes.

Discrete Markov chains whose state space is partitioned into disjoint classes which are periodically (with discrete period) visited by the chain are called periodic Markov chains [12]. By analogy, we refer to the cell-volume process also as periodic. Periodicity has consequences for the large-time behaviour of a process. Large-time behaviour of aperiodic processes is typically given by a steady-state distribution. Contrastingly, periodic processes retain the dependence of the phase even in the large-time limit.

In the next two sections, we will characterise the large-time behaviour of the periodic cell-volume process using first the single-cell and then the population approach.

## 5 Large-time single-cell behaviour

In the previous section we showed that the log-volume *u*(*t*) of a cell at time *t* is constrained to the set of values *n* + *φ*(*t*), where *n* is an integer the phase *φ*(*t*) is a function of time *t* (and of initial data). The probabilities

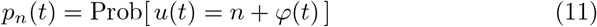

have a discontinuity at any time *t* for which at which *φ*(*t*) has a discontinuity (cf. Figure 1, bottom panel). We then have a consistency condition

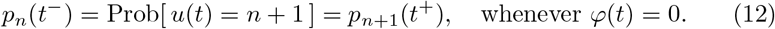

Away from the discontinuities, the probabilities (11) satisfy a system of balance equations

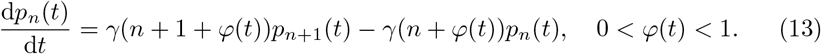

Integrating the system (13) forward in time, one obtains the probabilities *p_n_*(*t*) until a discontinuity in *φ*(*t*) is encountered. The consistency condition (12) needs to be applied at times of discontinuity to calculate from the probabilities right before the discontinuity their values right after the discontinuity. The integration of the system (13) can then be restarted with the post-discontinuity values.

The system (13) comprises an infinite number of coupled linear ordinary differential equations with non-constant coefficients. In general, the system (13) can be solved by truncating to a finite number of equations and using a numerical solver. In the specific case of a threshold dependence (7) of the division rate on the log-volume, we will be able to find an explicit large-time solution to (13) subject to (12).

As time progresses, the log-volume distribution becomes independent of the specifics of the initial condition and depend on time only via the phase *φ*. Let us denote this distribution by *π_n_*(*φ*). It satisfies a system of balance equations

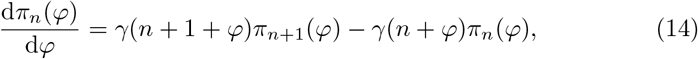

which looks similar to (13), differing in that the independent variable is now the phase *φ*, which is restricted to the range 0 ≤ *φ* ≤ 1. The consistency condition (12) translates into

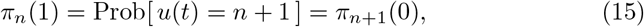

which provide a set of boundary conditions for the system (14). For threshold-type dependence (7) of division rate on log-volume, we have *γ*(*n* + *ϕ*) = *α*(1 – δ_*n*,0_), so that the system (14) simplifies to

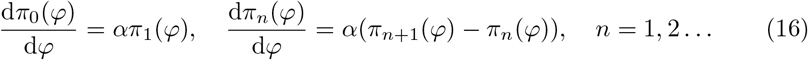

We look for a solution to (16) subject to the boundary conditions (15) in the form of an exponential

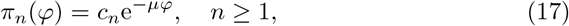

where *μ* is an eigenvalue and *c_n_* are eigenvector components. Inserting the ansatz (17) into (16) we find

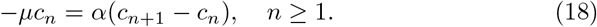

Substituting the ansatz (17) into (15) we find that

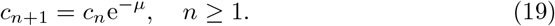

Substituting (19) into (18) and simplifying yields the characteristic equation

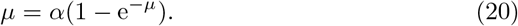

Elementary analysis shows that the characteristic equation (20) has a positive solution *μ* only if *α* > 1. Furthermore, it is a unique positive solution and lies in the interval *α* – 1 < *μ* < *α*. From now on we require that *α* > 1 holds and we take for *μ* the unique positive solution to the characteristic equation (20). The condition *α* > 1 guarantees the (post-threshold) dominance of division over growth, which is critical for the maintenance of cell volume homeostasis.

The recursive relation (19) implies that *c_n_* = *c*_0_e^−*μn*^, which, if substituted into (17), yields

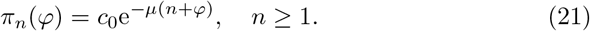

Positivity of *μ* guarantees that *π_n_*(*φ*) has a finite ℓ^1^ norm and can be normalised into a probability distribution. Integrating the first equation in (16) subject to *π*_0_(0) = 0 leads to

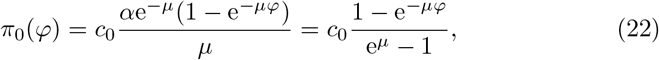

in which the second equality is due to (20). The normalisation constant c_0_ can be determined from the relation

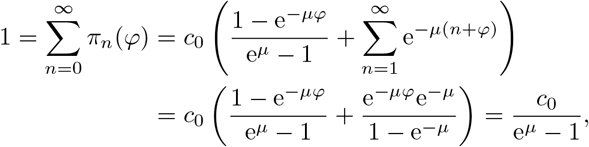

from which

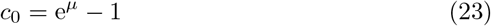

follows. Inserting (23) into (21) and (22) finalises our analysis.

In summary, we approximate the probability *p_n_*(*t*) that the cell’s log-volume is equal to *n* + *φ*(*t*) in the large-time regime by a phase-dependent distribution

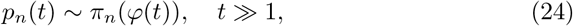

where *π_n_*(*φ*) is given explicitly by

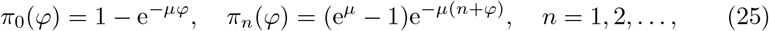

and *μ* is the unique positive solution to the transcendental characteristic equation (20), which exists provided that *a* > 1.

## 6 Large-time population behaviour

Assume that at the initial time *t*_0_ the population consisted of a single mother cell with log-volume *u*_0_. Algorithm 2 ensures that at *t* > *t*_0_ the log-volumes of its progeny are contained in a list 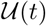. Lists differ from sets in that they can contain the same element multiple times. Due to periodicity of the cell-volume process, 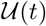 can only contain elements with values *n* + *φ*(*t*), where *n* is an integer and *φ*(*t*) is the phase as defined by (10). Define by *f_n_*(*t*) the number of times a particular value *n* + *φ*(*t*) is present in the list. The consistency condition

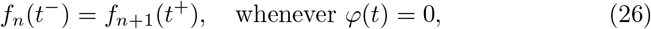

holds at times of discontinuity of *φ*(*t*). Provided that the numbers *f_n_*(*t*) are sufficiently large, we can treat them as continuous quantities that satisfy a population balance equation

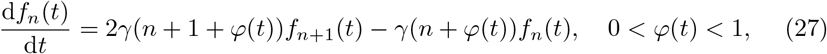

away from the times of discontinuity of *φ*(*t*). The population balance equation (27) differs from the probability balance equation (13) in the factor 2 multiplying the first term on the right-hand side of (27). It is easy to verify that *f_n_*(*t*) = 2^−*n*^*p_n_*(*t*), where *p_n_*(*t*) is a solution to the probability balance equation (13), is in fact a solution to the population balance equation (27). It does not however satisfy the consistency condition (26). In order to satisfy the consistency condition, we modify the solution to

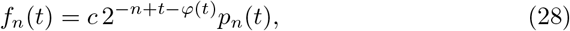

where *c* is a tunable constant. Since *t* – *φ*(*t*) is constant in any interval in which *φ*(*t*) is continuous, the function (28) is a constant multiple of 2^−*n*^*p_n_*(*t*) in any such interval, and as such satisfies the population balance equation (27). Further, it is easy to verify that the consistency condition (26) is met by (28). In the regime of large times, we can approximate *p_n_*(*t*) by *π_n_* (*φ*(*t*)) to obtain

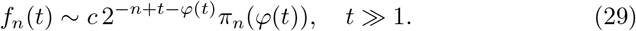

The total number of progeny at time *t* is given by

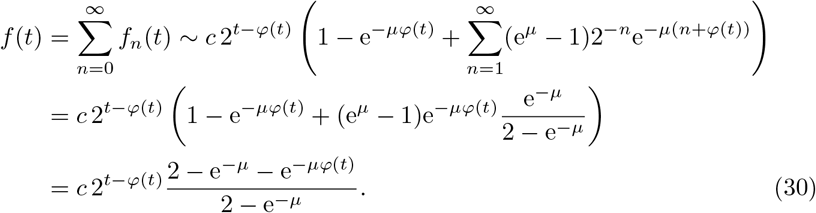

Finally,

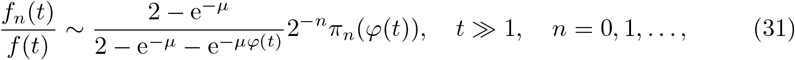

gives the proportion of cells which have log-volume *n* + *φ*(*t*) at a large time *t*.

## 7 Results

In this Section we use the simulation and analytic methods presented in the previous Sections to examine the dynamical behaviour of our model for cell-volume growth.

Figure 2 shows the cell count, on a logarithmic scale, as function of time measured in units of volume doubling time. Although the overall trend is characterised by an exponential increase, the cell count is nevertheless subject to periodically recurring cycles of fast growth alternating with slow growth. The cyclic behaviour is sustained even at large times. The simulation-based cell count (blue lines) exhibits low-copy-number noise at earlier times. The analysis-based cell count (orange lines) faithfully reproduces the large-time cyclic behaviour of the results of simulation. The four panels of Figure 2 differ in the choice of volume-dependent division rate *γ*(*V*). The value specified within the figure panels gives the division rate *α* that applies once a volume threshold has been crossed (cf. Equation (3)). For large values of the division rate *α* (bottom right panel in particular), the control of cell volume becomes near deterministic: division initiates almost immediately (i.e. with a very high rate) after the critical volume threshold is reached. We observe that in the near-deterministic regime of cell-volume control, the growth dynamics assumes a step-like pattern. Away from the deterministic regime, i.e. for lower values of the post-threshold division rate, the cycles of fast and slow growth are less pronounced.

**Fig. 2.**
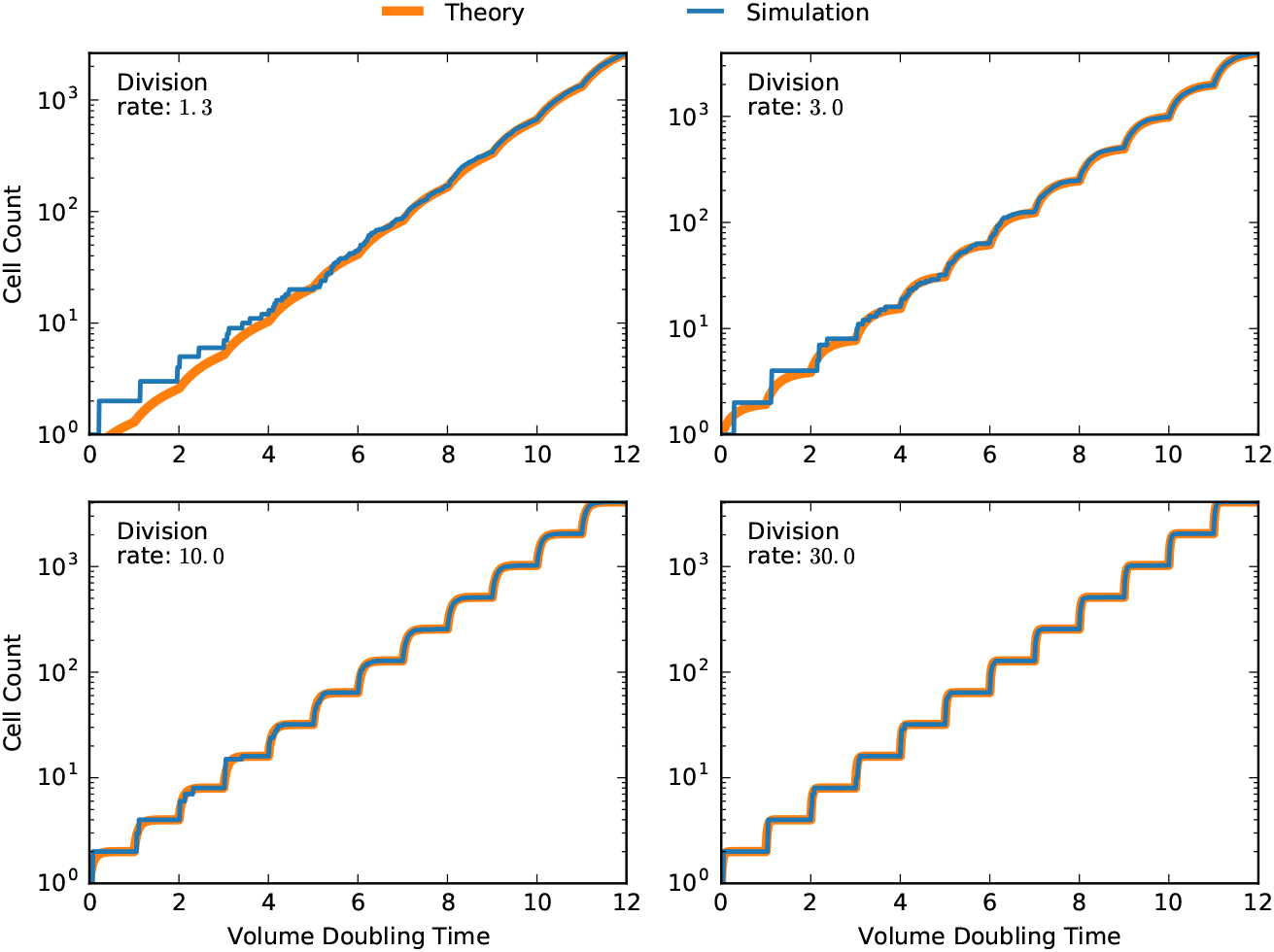
Size of the cell population derived from an individual mother cell as function of time measured in units of volume doubling time. Panels differ in the choice of the division rate *α* that applies after the critical cell volume has been reached by a growing cell. Results of stochastic simulation by Algorithm 2 (blue lines) cross-validate the theoretical prediction (30) (orange lines). The log-volume of the mother cell at initial time (here *t*_0_ = 0) is set to the critical value of *u*_0_ = 1 in all examples. The undetermined constant c in the theoretical result (30) has been chosen so as to perfectly fit the simulation result at the last timepoint (here *t_m_* = 12). The cell count was obtained by counting the total number of elements in the lists 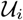 returned by Algorithm 2.

In the following computational experiment, we let a colony of cells, derived from a single progenitor, grow until it counts in thousand individuals, and then study in detail a single ensuing period of cyclic growth. The top panel in Figure 3 shows the dependence of the cell count on phase, which is consistent with the time dependence of cell count reported in Figure 2. In the remaining panels of Figure 3, blue-coloured bars represent the distributions of log-volume in the cell population at different phases of the period. At any phase of the cycle, the log-volume distribution is discrete (cf. Section 4). The support of the distribution, which is indicated by vertical dotted lines in the panels of Figure 3, travels to the right as phase increases. At the end of the period, the support of the distribution, as well as the distribution itself, returns to where it started at the beginning of the cycle. The theoretical proportions (31) (bars of lighter shade of blue) are in a good agreement with the results of simulation by Algorithm 2 (bars of darker shade of blue).

**Fig. 3.**
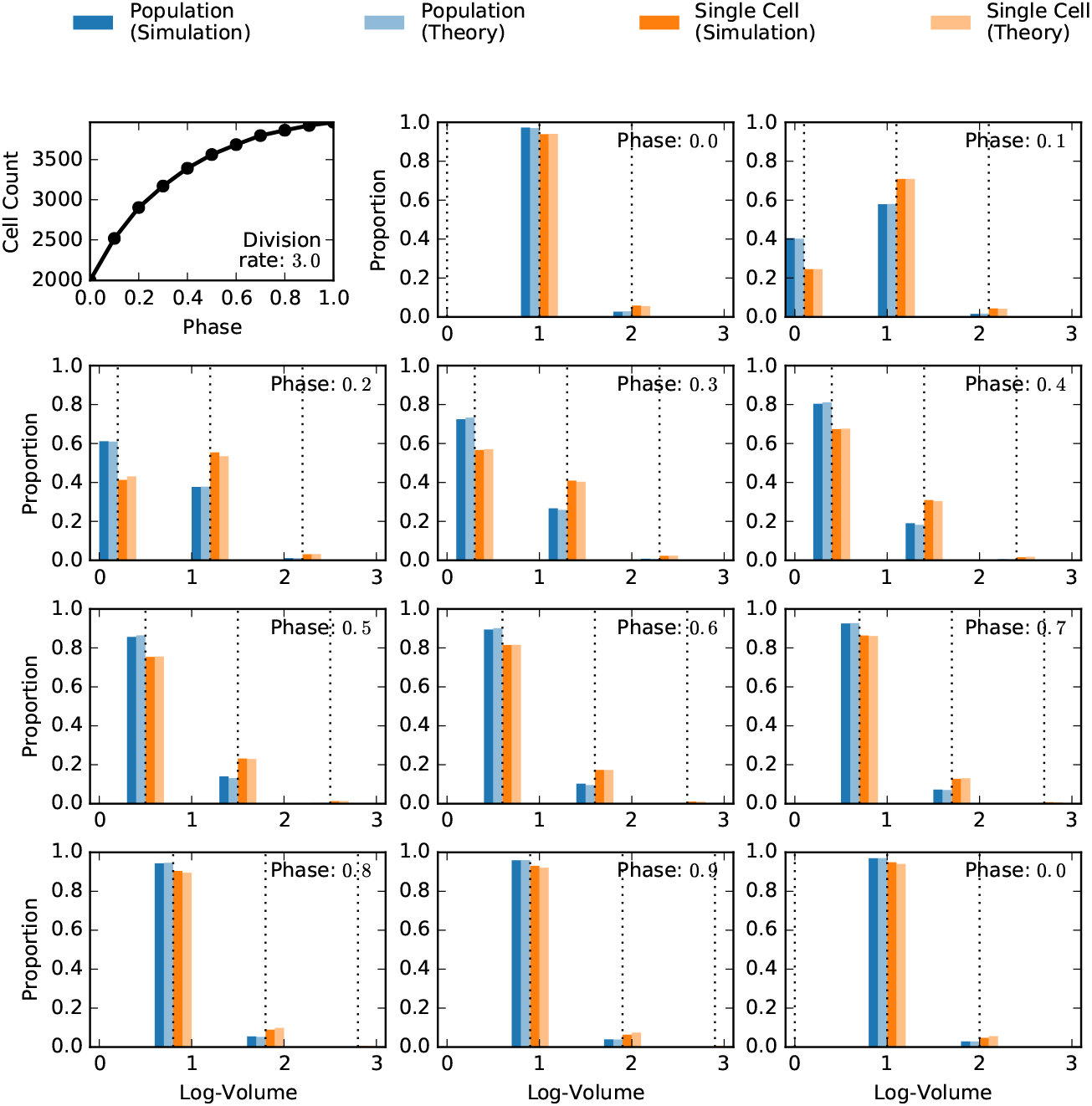
Discrete log-volume distributions at different phases of the growth cycle for post-threshold division rate set to *α* = 3.

In order to compare the population and single-cell versions of the model, we juxtapose the proportion of cell population with a particular log-volume (Figure blue bars) to the probability of observing the log-volume within a single-cell lineage (Figure 3, orange bars). The single-cell probabilities were estimated from an ensemble of 4096 independent sample paths generated by Algorithm 1 (Figure 3, bars of darker shade of orange). In each single-cell simulation, we used the same initial condition as in the population simulation; we also skipped the same amount of time, before analysing a single period of cyclic behaviour, as in the population simulation. The simulation-based estimates, both single-cell and population-wide versions, are cross-validated with the theoretical probability values (25) (Figure 3, bars of lighter shade of orange). Comparing the results of the population and single-cell versions of the model, we observe that the single-cell approach tends to underestimate the proportion of small cells and overestimate the proportion of big cells in the phases of fast growth (Figure 3, phases 0.1 to 0.7, say). In the later phases of slow growth, the two approaches lead to the same values of log-volume. These observations are yet more apparent in Figure 4 in which the division rate is set to a higher value *α* =10. The differences in the two approaches are noticeable only at the early phases of very fast growth (Figure 4, phases 0.1 to 0.3). In the later phases, the two approaches give the same prediction.

**Fig. 4.**
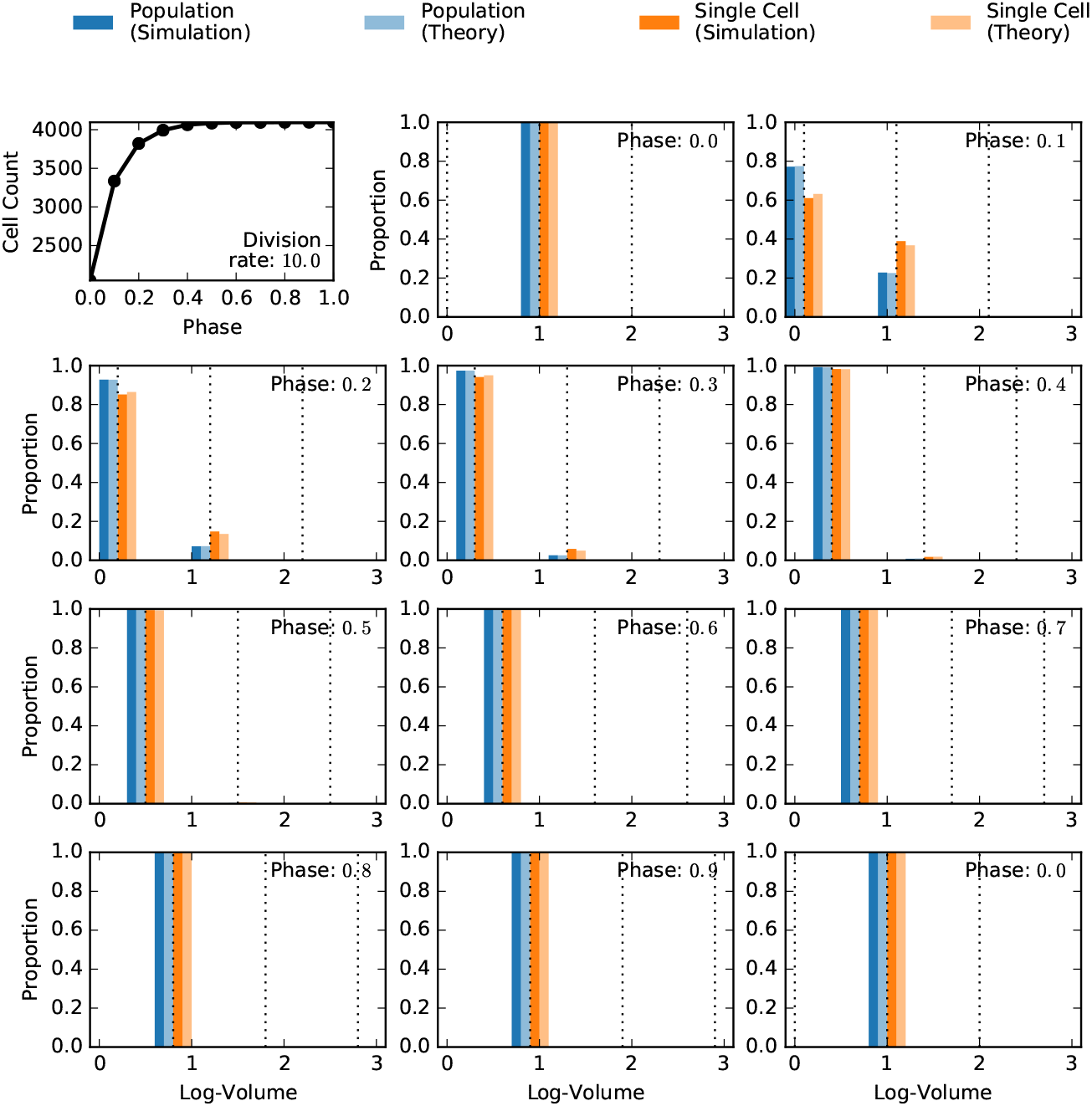
Discrete log-volume distributions at different phases of the growth cycle for post-threshold division rate set to *α* = 10.

## 8 Conclusions

We considered a model for the growth of cellular volume in which cells grow exponentially and divide with a volume-dependent rate, resulting in two daughter cells each inheriting half of their mother’s volume. Two versions of the model are systematically compared: in the first version, a single cell lineage is described by a stochastic, Markovian, model; in the second, an exponentially growing cell population is followed. We constructed simulation algorithms for both model versions which are exact in the sense that they require no numerical discretisation technique to sample the underlying stochastic process [8].

The model, in either of the two formulations, is shown to sustain cyclic behaviour with alternating phases of slow and fast growth. The phases of fast growth occur when most cells are large enough to divide, whereas the phases of slow growth take place when most cells are too small to divide. The cyclic behaviour, resulting from the periodicity of the stochastic process, is intriguing as it suggests possible biases from specific experimental designs (e.g. choice of measurement times). Notably, periodicity is a consequence of a fully symmetric division, and even small amounts of asymmetry in a more general model situations are expected to eventually break periodicity.

Our computational analysis suggests that the population-based approach leads to greater proportions of small cells and smaller proportions/probabilities of large cells in the fast-growth phases than the single-cell approach. This observation is consistent with the fact that the population approach includes twice as many daughter cells than mother cells do in comparison with the single-cell approach. Additionally, our results provide a quantitative evaluation of this effect and tractable analytic solutions valid in the large time asymptotic regime.

Thus, our results provide insights into the dynamics of the process of cell growth and suggest commonalities as well as differences between single-cell Markovian modelling and whole-population simulation. We expect that the methods explored in this paper can be applicable in related and more complex descriptions of cell growth and division.

## Bibliography

[1] Alberts, B., Johnson, A., Lewis, J., Raff, M., Roberts, K., Walter, P.: Molecular biology of the cell. Garland Science, New York (2002)

[2] Amir, A.: Cell size regulation in bacteria. Phys. Rev. Lett. 112(20), 208102 (2014)

[3] Antunes, D., Singh, A.: Quantifying gene expression variability arising from randomness in cell division times. J. Math. Biol. 71(2), 437–463 (2015)

[4] Bell, G.I., Anderson, E.C.: Cell growth and division: I. a mathematical model with applications to cell volume distributions in mammalian suspension cultures. Biophys. J. 7(4), 329 (1967)

[5] Bernard, E., Doumic, M., Gabriel, P.: Cyclic asymptotic behaviour of a population reproducing by fission into two equal parts. arXiv preprint arXiv:1609.03846 (2018)

[6] Davis, M.: Piecewise-deterministic markov processes: A general class of non-diffusion stochastic models. J. R. Stat. Soc. B 46, 353–388 (1984)

[7] Diekmann, O., Heijmans, H.J., Thieme, H.R.: On the stability of the cell size distribution. J. Math. Biol. 19(2), 227–248 (1984)

[8] Gillespie, D.: Exact stochastic simulation of coupled chemical reactions. J. Phys. Chem. 81, 2340–61 (1977)

[9] Hannsgen, K.B., Tyson, J.J., Watson, L.T.: Steady-state size distributions in probabilistic models of the cell division cycle. SIAM J. Appl. Math. 45(4), 523–540 (1985)

[10] Kretzschmar, K., Watt, F.M.: Lineage tracing. Cell. 148, 33–45 (2012)

[11] Modi, S., Vargas-Garcia, C.A., Ghusinga, K.R., Singh, A.: Analysis of noise mechanisms in cell-size control. Biophys. J. 112(11), 2408–2418 (2017)

[12] Norris, J.R.: Markov chains. Cambridge Univ Press, Cambridge, UK (1998)

[13] Perthame, B.: Transport equations in biology. Springer Science & Business Media, Berlin/Heidelberg (2006)

[14] Robert, L., Hoffmann, M., Krell, N., Aymerich, S., Robert, J., Doumic, M.: Division in escherichia coli is triggered by a size-sensing rather than a timing mechanism. BMC biology. 12(1), 17 (2014)

[15] Schuss, Z.: Theory and applications of stochastic processes: an analytical approach. Springer Science & Business Media, Berlin/Heidelberg (2009)

[16] Taheri-Araghi, S., Bradde, S., Sauls, J.T., Hill, N.S., Levin, P.A., Paulsson, J., Vergassola, M., Jun, S.: Cell-size control and homeostasis in bacteria. Curr. Biol. 25, 385–391 (2015)

[17] Thomas, P.: Analysis of cell size homeostasis at the single-cell and population level. Front. Phys. 6, 64 (2018)

[18] Vargas-Garcia, C.A., Ghusinga, K.R., Singh, A.: Cell size control and gene expression homeostasis in single-cells. Curr. Opin. Syst. Biol. 8, 109–116 (2018)

[19] Vargas-Garcia, C.A., Soltani, M., Singh, A.: Conditions for cell size homeostasis: a stochastic hybrid system approach. IEEE Life Sci. Lett. 2(4), 47–50 (2016)

